# Doxorubicin induces cardiotoxicity in a pluripotent stem cell model of aggressive B cell lymphoma cancer patients

**DOI:** 10.1101/2020.04.15.042424

**Authors:** Luis Peter Haupt, Andreas Maus, Malte Tiburcy, Steffen Köhne, Wiebke Maurer, Rewati Tappu, Jan Haas, Yun Li, Andre Sasse, Celio C. X. Santos, Ralf Dressel, L. Wojnowski, Gertrude Bunt, Ajay M. Shah, Benjamin Meder, Samuel Sossalla, Bernd Wollnik, Gerd Hasenfuß, Katrin Streckfuß-Bömeke

## Abstract

Cancer therapies have been shown to induce cardiovascular complications. The aims of this study were to establish an *in vitro* induced pluripotent stem cell model (iPSC) of anthracycline-induced cardiotoxicity (ACT) from patients with an aggressive form of cancer.

ACT-iPSC-CM generated from individuals with CD20^+^ B-cell lymphoma cancer who had received high doses of DOX and suffered cardiac dysfunction were observed to be persistently more susceptible to DOX toxicity compared to control-iPSC-CM. ACT-iPSC-CM exhibited increased DOX-dependent disorganized myofilament structure and cell death, as well as higher reactive oxygen species (ROS) compared to controls. Importantly, analysis of engineered heart muscle (EHM) from ACT-iPSC-CM showed an impaired DOX-dependent mechanical functionality. Transcriptome profiles of EHM are in line with a disturbed adjustment to DOX-dependent alteration of Ca^2+^ homeostasis in ACT-iPSC-CM. Furthermore, genetic variants in different cardiac key regulators were uncovered.

In conclusion, we developed the first human iPSC-CM and EHM model of DOX-induced cardiac dysfunction in patients with B-cell lymphoma. Our results suggest that DOX-related stress resulted in decreased contractile activity and finally in heart failure in ACT patients.

**Brief summary:** Development of the first human iPSC-CM model of DOX-induced cardiac dysfunction in patients with aggressive B cell lymphoma and high-dose DOX treatment.

## Introduction

Anthracycline-induced cardiotoxicity (ACT) was first described in 1971(1). Although it has been known for decades that the anthracycline drug doxorubicin (DOX) can trigger cardiotoxicity, its outstanding efficacy against a broad range of solid and hematopoietic cancers often overrides considerations of its risk for the clinical application. Currently it is administered to 32% of breast cancer patients(2), 57-70% of elderly lymphoma patients(3, 4), and 50-60% of childhood cancer patients(5). After it became evident that the risk of ACT rises with increasing doses of DOX, the lifelong cumulative dose of DOX was set at 500 mg/m^2,^ (6, 7). Nevertheless, more recent studies suggest that the incidence of ACT is still at 5-9%, with up to 18% of DOX-treated patients showing subclinical symptoms (8-10). The side effects of cardiotoxicity include disturbance in ventricular de/-repolarization, arrhythmia, decrease in left ventricular ejection fraction (LVEF), and fractional shortening (FS) often leading to dilative heart failure (HF) and enhanced mortality (11).

Although ACT causes lethal congestive heart failure (HF), no early detection or effective treatment methods are yet available(12). Development of ACT is highly inter-individually variable, and is thus not possible to accurately predict its occurrence. Why only some patients develop ACT, and exactly why it developed in these and not others, has been the subject of intense study. Recently, clinical study cohorts were investigated for an association of gene variants with ACT (13-15) including the ‘rituximab with CHOP over age 60 years’ (RICOVER60) trial (16) and the NHL-B1/B2 study.

The pathophysiology of ACT is described as multifactorial and the exact causal molecular mechanisms of DOX-induced cardiotoxicity remain elusive. The main cardiotoxic effects of DOX observed so far include elevated levels of reactive oxygen species (ROS), cardiomyocyte death, mitochondrial dysfunction, topoisomerase II β poisoning, induction of autophagy and defective calcium homeostasis (17-19). However, the mechanism through which DOX damages the heart and causes disease progression over multiple decades is still poorly understood.

Generation of ROS by so-called redox cycling in cardiomyocytes can cause lipid peroxidation, DNA damage, and mitochondrial dysregulation. Mitochondria are the major site of DOX-dependent ROS production due to the localization of redox cycling enzymes as NADH oxidases. It is likely that DOX interferes with the electron transport chain, disrupts mitochondrial membranes, and further increases mitochondrial ROS production(20, 21). Dysregulated redox signaling is known to be connected to cardiac diseases (22-24). Other proposed mechanisms include inhibition of topoisomerase II by DOX’ intercalation into DNA, leading to DNA damage-induced cellular stress and cell death in cardiomyocytes (25-27) which in turn result in loss of functional cardiomyocytes and heart injury (19). Finally, DOX induces disruption of Ca^2+^ homeostasis in CM by increasing Ca^2+^ release from the sarcoplasmic reticulum (SR) by higher opening probabilities of ryanodine receptor 2 (RYR2) (18, 28, 29). On the other hand, expression and activity of the sarco/endoplasmic reticulum Ca^2+^ATPase (SERCA) is reduced by DOX (18, 30), leading to decreased Ca^2+^ transport into the SR and cytoplasmic Ca^2+^ overload. This results in sarcomeric disarray and a reduction in the heart’s contractile force, and may contribute to generation of arrhythmia (18, 31).

Not only cardiomyocytes but also cardiac fibroblasts (cFB) are key players in myocardial pathology. cFB are considered the predominant source of the extracellular matrix and the hallmark of fibrosis (32). During cardiac stress, cFB trans-differentiate into myofibroblasts, which perform cardiac extracellular remodeling and gain contractile activity due to alpha smooth muscle (α-SMA) expression(33, 34). An increase in the extracellular matrix (ECM) normally results in cardiac stiffness and diastolic dysfunction as well as impaired cardiac contraction (35, 36). CFB interact with CM through direct connections and paracrine signaling (37). However, less is known about the direct effects of DOX on human cFB from diseased ACT myocardium.

There has been a lack of human cardiomyocyte culture models from patients with ACT that could be used to help us better understand the molecular and cellular physiology of ACT. However, the generation of induced pluripotent stem cell-derived cardiomyocytes (iPSC-CM) has now made it possible to establish ACT patient-specific disease models. iPSC CM have been widely used to study hereditary and multifactorial cardiac conditions in vitro, including arrhythmic disorders or (stress) cardiomyopathies, with a correlation to predicted phenotypes (38, 39). Patient-specific stem cell models showed a predilection of breast cancer patients to low DOX-induced cardiotoxicity (40) or trastuzumab-induced cardiac dysfunction(41). A genetic predisposition has been suggested for ACT (13), but this is not yet established.

In the present study, we used iPSC-CM, cFB, and engineered heart muscle (EHM) generated from patients who received high dosages of DOX to investigate their contribution to the development of cardiac dysfunction. We show here that iPSC-CM, cFB and EHM generated from ACT patients are more susceptible to detrimental effects of DOX treatment than CM, cFB and EHM from control patients. Transcriptome analysis revealed the key role of an altered mRNA translational process for iPSC-CM in disease pathogenesis. The resulting inability of ACT-iPSC CM to adapt to acute DOX exposure as control iPSC-CM were able to do was demonstrated by the DOX-induced disruption of the Ca homeostasis. Control iPSC CM increased protein expression of Ca^2+^ transporting proteins in the sarcoplasmic reticulum as SERCA and RYR2, and thus avoided cytoplasmic Ca^2+^ overload and heart failure conditions. In contrast, ACT-iPSC-CM showed decreased SERCA protein but DOX-induced CamKIIδ activity of target phosphorylation of RYR2-S2814 and PLN-Thr17. These findings have made it possible for us to develop a simultaneous modulation of SERCA and CamKII activity which may serve as a novel therapeutic and protective strategy for patients with ACT.

## Results

### Characterization of human ACT myocardium

To investigate the importance and severity of ACT, we had the unique opportunity to analyze samples of human explanted myocardium from 5 patients with end-stage heart failure that developed after anthracycline chemotherapy, and compare these to samples of healthy non-failing (NF) controls (Table 1). Fibrosis was analyzed using Masson’s trichrome staining followed by quantification of fibrotic areas in relation to myocardium. In the ACT patients, we found massive fibrosis in 24.4%, a rate significantly higher compared to healthy controls (8.5%) (Figure 1A, B). Accordingly, there was a significantly increased expression of fibrotic markers such as connective tissue growth factor (CTGF) and matrix metalloproteinase 9 (MMP9) in the ACT patient samples (Figure 1C). In addition, we examined the expression of SERCA2 as a key player in excitation/contraction coupling and found it to be significantly downregulated on the mRNA and protein level in ACT patients (Fig. 1D). Redox stress-associated NADPH oxidase subunits as *NOX2, RAC1, RAC2, NCF2, and NCF4* were substantially downregulated in ACT tissue compared to NF, whereas NOX4 was upregulated in ACT myocardium compared to NF (Supplementary Figure 1A-c). These data confirmed that fibrosis is a significant feature of end-stage ACT myocardium, and that oxidative stress may play a key role in ACT development.

**Table 1:** Characteristics of end-stage heart failure ACT patients

| patient | LVEF [%] | Sex | Age at TX | Cancer type |
| --- | --- | --- | --- | --- |
| **ACT-4** | 20 | female | 65 | Mammary carcinoma |
| **ACT-5** | 10-15 | male | 54 | Ewing sarcoma |
| **ACT-6** | 20 | female | 53 | Wilms tumor |
| **ACT-7** | 20 | male | 16 | B cell non-Hodgkin gastric lymphoma |
| **ACT-8** | 15 | male | 43 | Unknown |

**Figure 1:**
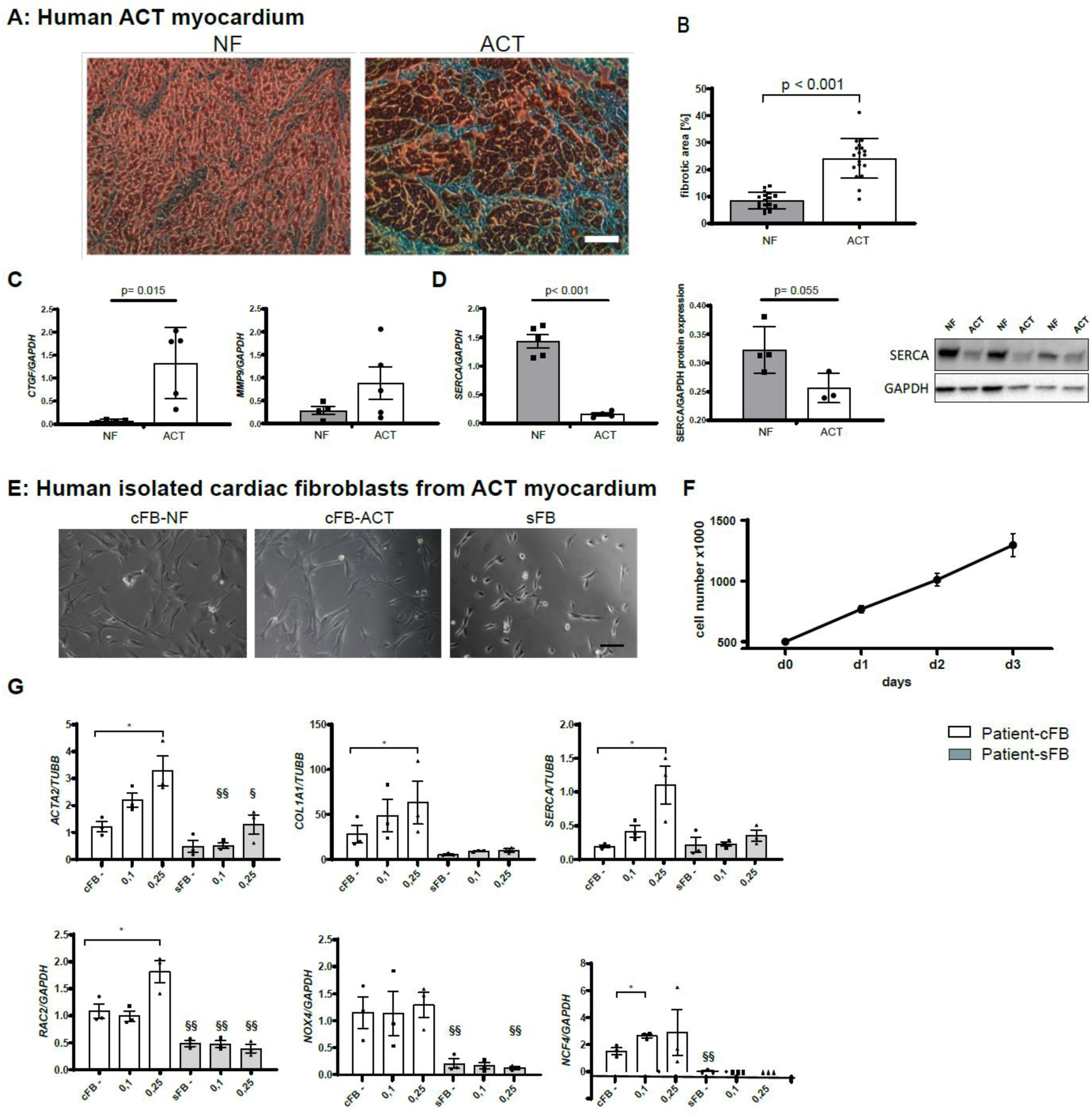
Analysis of human myocardium of ACT patients and isolated ACT cardiac fibroblasts. **(A)** Representative myocardium of non-failing donor (NF) and ACT patients (ACT) after trichrome staining. Scale bar: 100 µm. **(B)** Quantification of trichrome staining; NF: n=5; ACT: n=5 (biological replicates). 3 technical replicates from different tissue slices for each biological replicate were used. **(C)** mRNA expression of fibrosis-associated genes (*CTGF, MMP9*) in human ACT myocardium compared to NF by qPCR. **(D)** mRNA and protein expression of SERCA. Representative Western Blots for SERCA in NF (n=4) and ACT (n=3) myocardium are shown. GAPDH was used as an internal control. Statistical analysis was performed using Student’s t-test. p values: P >0.05 are defined as statistically significant. Bars indicate mean values ± SEM. **(E)** NF cardiac fibroblasts (cFB-NF), cardiac fibroblasts from the ACT patient (Patient cFB), and skin FB of ACT patient (Patient sFB) show typical FB morphology. Scale bar: 100 µm. **(F)** Proliferation capacity over three days of human cardiac fibroblasts from 5 end-stage heart failure patients. **(G)** Relative DOX-dependent mRNA expression of fibrosis- and Ca^2+^-associated genes and NADPH oxidase subunits by qRT-PCR for the genes *ACTA2, COL1A1, SERCA, NOX4, RAC2*, and *NCF4* in cFB from the ACT patient (Patient-cFB) and sFB (patient-sFB) under basal condition (-) and different DOX treatment concentrations (0.1 µM and 0.25 µM). n=3 biological replicates (samples from three different passages per cell type) for each measurement. Statistical analysis was performed with 1-way ANOVA analysis or Student’s t-test. * p < 0.05 in the cFB group and as § p < 0.05, §§ p < 0.01 between cFB and sFB. bars indicate mean values with± SEM.

### Characterization of isolated human cardiac fibroblasts from ACT myocardium

To investigate the hypothesis that cFB contribute together with CM to the development of ACT, cFB were isolated from the left ventricle of ACT human myocardium and characterized as to their origin. We were able to demonstrate that isolated cardiac cells have a characteristic FB morphology and a high proliferative capacity, and express typical FB genes and proteins (Figure 1E, Supplementary Figure 1D-E). DOX treatment of ACT-cFB and skin FB (sFB) revealed that DOX markedly increased the expression of *ACTA2* and *COL1A1*, indicating trans-differentiation into active ACT-cFB compared to mostly unchanged expression in sFB (Figure 1G). Interestingly, *SERCA* expression was enhanced upon DOX treatment in ACT-cFB, but not in sFB (Figure 1G). We observed that certain NADPH oxidase subunits such as *RAC2* and *NCF4* increased in expression after DOX, but there was little effect on *NOX4* in both ACT-cFB and sFB. Furthermore, expression levels were markedly decreased in sFB for all tested markers compared to ACT-cFB (Figure 1G). These results indicate that cFB isolated from human ACT myocardium exhibits an increased myofibroblastic stress response after DOX treatment.

### Recruitment of ACT patients for iPSC generation

We aimed to generate iPSC CM from ACT patients to perform a patient-specific analysis of the long-term DOX-effects in human CM. For this we recruited five patients from the ‘rituximab with CHOP over age 60 years’ RICOVER60 trial (www.clincaltrials.gov: NCT00052936). All of these patients suffered from diffuse large B cell lymphoma and had been treated with DOX as part of a CHOP-14 treatment (cyclophosphamide, doxorubicin, vincristine and prednisolone in two-week intervals). RICOVER60 trial participants were classified into ACT patients and controls based on Reichwagen et al., 2015, with symptoms such as reduced left ventricular ejection fraction (LVEF <45%), arrhythmia, or heart failure treatment for ACT (14). Based on these criteria, we recruited three patients for our study (referred to as ACT patients) who suffered from clinical cardiotoxicity (post-treatment LVEF: <45%) as well as two patients (controls) who did not experience clinical cardiotoxicity after chemotherapy (post-treatment LVEF: >55%) (Table 2).

**Table 2:** Characteristics of ACT patients and controls from the RICOVER60 trial used for iPSC generation

| **patient** | **Sex** | **Age in 2007** | **Median DOX**<br>**treatment [mg /m<sup>2</sup>]** | **ACT** |
| --- | --- | --- | --- | --- |
| **ACT-1** | male | 69 | 309 | chronic |
| **ACT-2** | male | 71 | 309 | chronic |
| **ACT-3** | female | 66 | 309 | chronic |
| **Control 1** | male | 66 | 318 | no |
| **Control 2** | male | 68 | 318 | no |

### Generation and characterization of iPSC-derived cardiomyocytes

We generated integration-free iPSCs from dermal fibroblasts from three ACT patients and two controls from the RICOVER60 trial. Two independent iPSC cell lines were generated per patient and analyzed for their pluripotency. All generated iPSCs maintained full pluripotency and spontaneous in vitro and in vivo differentiation capacity (Supplementary Figure 2A-C). Both ACT- and control-iPSCs were differentiated into CM using standardized WNT modulation (42) and metabolic selection (43). Individual batches were tested for homogeneity with staining for the cardiac specific marker cTNT and subsequent analysis with flow cytometry at day 60 of differentiation. Our differentiations consisted of more than 95% cTNT-positive cells (Supplementary Figure 2D). IPSC CM expressed mRNA for α-actinin, cTNT, α-MHC and β-MHC (Supplementary Figure 2E), whereas expression of these genes was much lower in undifferentiated iPSCs.

### DOX resorption and retention in iPSC CM of ACT patients and controls

Differences in the pharmacokinetics between patients can result in vastly different efficacy or side effects. We therefore treated the ACT CM with increasing DOX concentrations or durations based on pharmacokinetic characteristics of DOX in humans (44), and analyzed the intracellular DOX levels in ACT CM in comparison to control CM. Both ACT CM and control CM showed a positive correlation between DOX treatment concentration and intracellular DOX levels, with consistently higher intracellular DOX levels in ACT-iPSC CM (Figure 2A). Time-dependent DOX resorption experiments demonstrated increased intracellular DOX concentrations with peak values after 48 h of treatment (Figure 2B). Since chronic ACT can manifest itself many years or even decades after cancer treatment (45), we explored the hypothesis that remaining small amounts of DOX continuously cause damage in CM. iPSC CM were treated with DOX for 24 h and recovered for an additional three or seven days, respectively. DOX levels were significantly decreased to 25% after 3 d of recovery in control iPSC CM, whereas a similar decrease to 17.3% of intracellular DOX was demonstrated in ACT iPSC CM after at least seven days of recovery (Figure 2C). These results show that DOX did not remain in CM for extended periods and that DOX remains longer in iPSC CM from ACT patients compared to controls. Of note, on the basis of this data as well as prior reports (40), we primarily selected the time point of 24 h (for most experiments) and used DOX at concentrations in the range of 0.1-5 µM for all further experiments in iPSC-CM, cFB and EHM.

**Figure 2:**
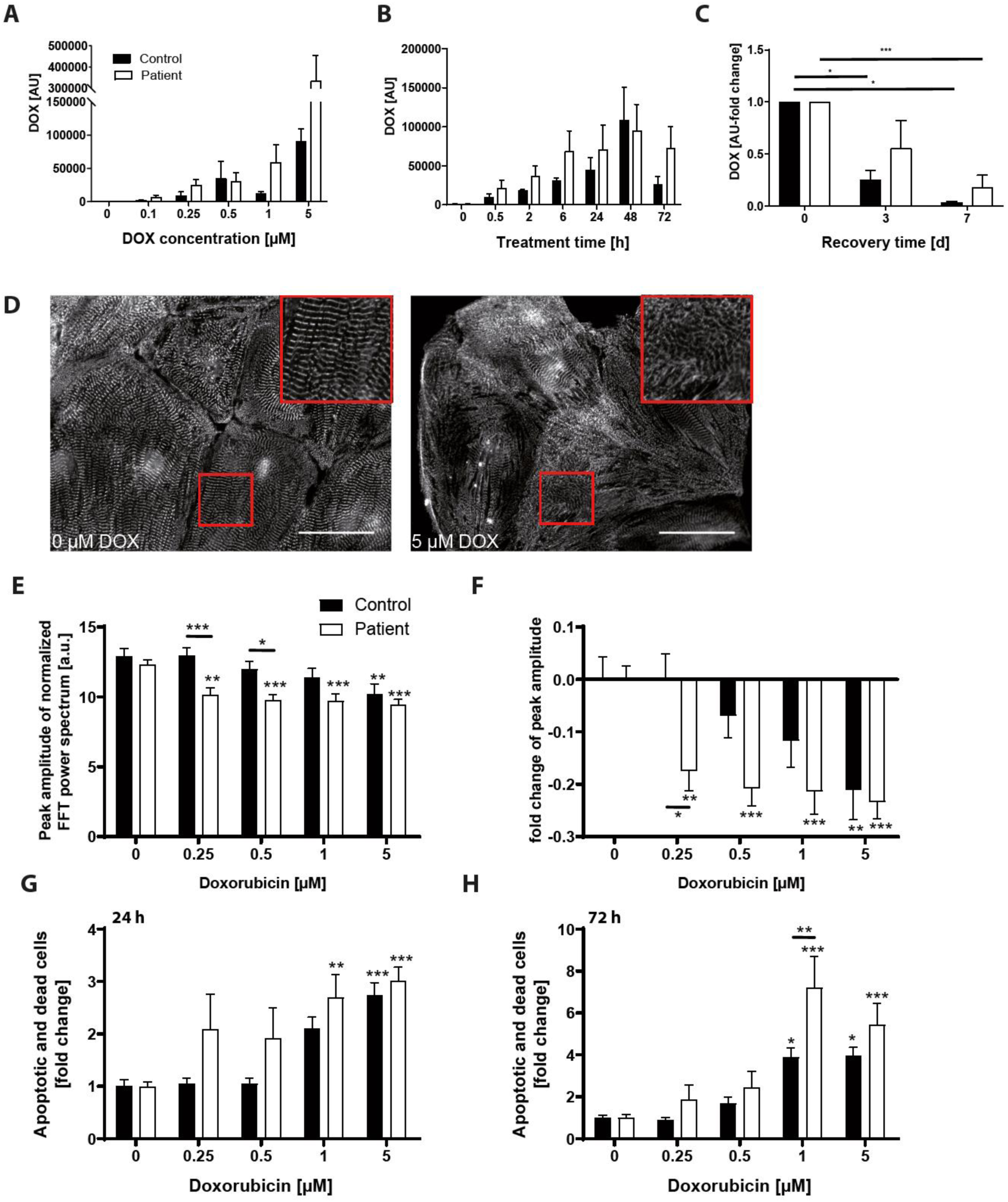
Assessment of in vitro doxorubicin-induced cardiotoxicity in patient-specific iPSC-CM. (A-C) DOX resorption and retention in iPSC-CM. Intracellular DOX levels were investigated with HPLC in regard to DOX concentration (24 h treatment) (A), treatment time (1 µM DOX) (B), and recovery (0, 3, 7 days) after treatment (1 µM DOX for 24 h) (C). ACT patient: n= 4-5 differentiations from 2 cell lines from 2 patients. Control: n= 3-4 differentiations from 2 cell lines from 1 patient. **(D-F)** Effect of doxorubicin on sarcomeric regularity in iPSC-CM **(D)** Immunofluorescence staining visualized α-actinin protein expression and sarcomeric organization. DOX treatment impaired sarcomeric regularity. Scale bars: 50 μm. (**E, F**) Quantification of DOX-treated (24 h) sarcomeric regularity using Fast Fourier Transform algorithm. Sample numbers: 60-62 control-iPSC-CM from 4 differentiations, 85-91 patient iPSC-CM from 6 differentiations. (**G, H**) Effect of doxorubicin on iPSC-CM viability. Annexin V/PI apoptosis tests showed that the number of apoptotic and dead cells rose with increasing DOX levels after 24 h (**G**) and 72 h (**H**). 10 control CM differentiations, 10 patient CM differentiations. Statistical analysis was performed using 2-way ANOVA analysis and Sidak’s or Dunnett’s multiple comparison, or mixed-effects analysis. Bars indicate mean values with SEM. * p < 0.05, ** p < 0.01, *** p < 0.001.* above single bars indicate statistical significances to untreated (0 µM DOX) conditions of the same group.

### DOX-induced cardiotoxicity is increased in ACT-iPSC CM

Direct myofibril damage can be visualized by α-actinin staining and is a hallmark of doxorubicin cardiotoxicity (46). To assess the sarcomeric integrity after treatment with DOX, we analyzed the sarcomeric structures in ACT and control iPSC CM. The sarcomeric regularity was visibly impaired after treatment with 5 µM DOX for 24 h (Figure 2D). Quantification using Fast Fourier Transformation (FFT) and radial integration confirmed the dose-dependent decrease of sarcomeric regularity in both groups (Figure 2E). Using the clinically relevant concentration of 0.25 µM DOX, the impairment of sarcomeric regularity was significantly greater in ACT CM compared to control CM (Figure 2D-F).

To analyze programmed cell death, we performed flow cytometry of iPSC CM that were positively stained for both annexin V and propidium iodide (PI). We found a dose-dependent relative increase in both apoptotic and dead cells after DOX incubation for 24 h and 72 h in both groups, suggesting that programmed cell death seems to be a principal mechanism of DOX-induced cell loss (Figure 2 G, H). All DOX concentrations tested resulted in increased amounts of apoptotic and dead cells in the ACT patient group compared to controls (Figure 2G, H). A significant increase in apoptotic and dead cells was detected in ACT CM compared to control CM at a DOX concentration of 1 μM for 72 h (Figure 2H). These data suggest that DOX-treated iPSC CM recapitulate ACT disease phenotypes such as sarcomeric organizational damage and programmed cell death, as is commonly described in ACT patients.

### DOX-dependent ROS production is enhanced in ACT iPSC CM and ACT cFB compared to control

Since the increased generation of reactive oxygen species (ROS) is described as one of the key mechanisms underlying the cardiotoxic effects of DOX, we aimed to assess the amount of ROS in ACT iPSC CM compared to controls by measuring extracellular H2O2 with Amplex Red. After 24 h of DOX treatment, H2O2 increased DOX-dependently with significantly higher H2O2 at 0.5 µM DOX in ACT iPSC CM compared to control cells (Figure 3A). Higher DOX concentrations do not influence H2O2 levels in ACT or control iPSC CM. We addressed the question as to whether the observed DOX-induced changes in H2O2 production occurred only during or immediately after the treatment of the cells, and analyzed the H2O2 level 7, 14 and 21 days after one-time DOX treatment. We found a 2-fold increase in H2O2 directly after the 24 h DOX treatment (0 days after DOX treatment) and similar H2O2 amounts 14 and 21 d after DOX treatment in both groups (Figure 3B). Interestingly, a significant 3-4-fold increase in H2O2 was found 7 d after DOX treatment in control and ACT iPSC CM (Figure 3B). These findings suggest that chronic changes were induced by single DOX applications because almost no DOX was detectable in the cells at the same time points (7 d after DOX treatment) (Figure 2C). Since we found a fibrotic phenotype of ACT myocardium, we analyzed DOX-induced ROS production in ACT cFB, ACT sFB and healthy control cFB. We found a DOX-dependent H2O2 increase in all tested FBs with significantly higher H2O2 amounts at all measured DOX concentrations in ACT cFB compared to healthy cFB or patient sFB (Figure 3C). Taken together, both ACT-iPSC CM and ACT cFB showed dose-dependent increases in extracellular H2O2 compared to healthy controls, suggesting a contribution of both cell types to the development of ACT.

**Figure 3.**
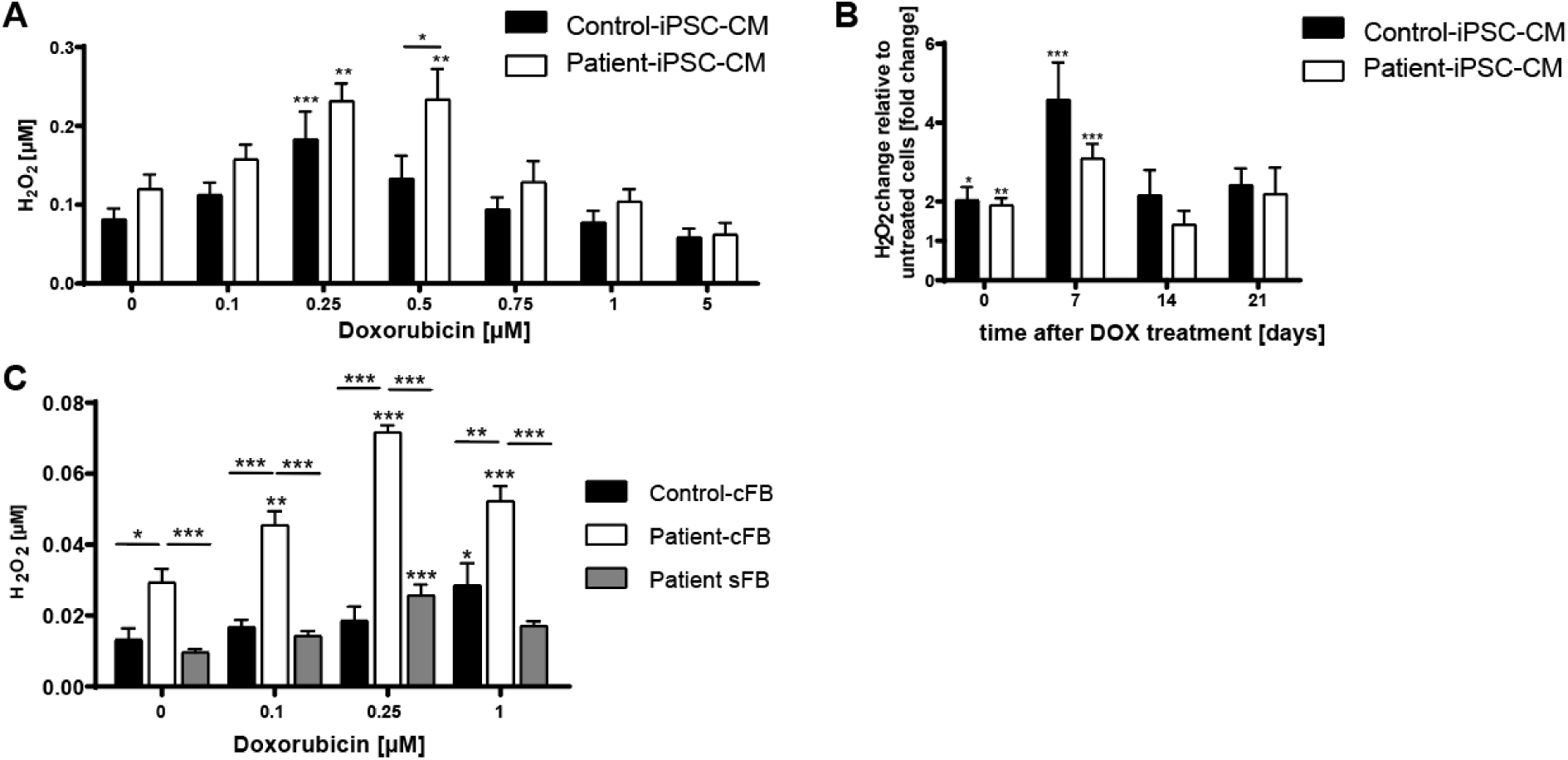
DOX-dependent ROS production in iPSC-CM and cFB. (**A**) The amount of H2O2 in the supernatant of iPSC-CM was measured with the Amplex Red assay after 24 h DOX treatment. (**B**) Relative changes in H2O2 amount in iPSC-CM of ACT patient and control 0, 7, 14 and 21 days after single treatment with 0.25 µM DOX for 24 h. **(C)** H2O2 in the supernatant of ACT patient- and control cFB, and patient skin FB (patient-sFB) was measured with the Amplex Red assay after 24 h DOX treatment. Sample numbers: (A) 5-10 control-CM differentiations, 7-17 ACT patient CM differentiations. (B) 11 control-CM differentiations, 17 ACT patient CM differentiations. (C) n=3 biological replicates (samples of 3 different passages were measured of control cFB, patient cFB, and patient sFB. Statistical analysis was performed using 2-way ANOVA analysis and Sidak’s or Dunnett’s multiple comparison test. Data are shown as mean with SEM. * p < 0.05, ** p < 0.01, *** p < 0.001. .* above single bars indicate statistical significances to untreated (0 µM DOX) conditions of the same group.

### Decreased DOX-dependent contractile function in ACT patients

To investigate how the DOX-dependent alterations in sarcomeric integrity, ROS production, and apoptosis affect muscle function, we generated EHM from ACT or control iPSC CM. EHM from ACT-iPSC CM and control iPSC CM exhibited comparable spontaneous beating frequencies (control: 35.8 ± 3.5, ACT: 28.4 ± 2.9 bpm). DOX treatment induced an increase in beating frequency, which was significantly more pronounced in ACT EHM (Figure 4A) compared to controls. About 15% of generated EHM displayed irregular beating at basal conditions and DOX treatment caused the rate to increase to 40% in both groups (Supplementary Figure 3B). Furthermore, EHM from both groups showed similar maximal force of contraction at increasing Ca^2+^ concentrations (Figure 4B). Upon DOX treatment, the force of contraction was significantly decreased in the ACT patient group but not in the control group (Figure 4B, C, D). Despite the significant decrease in cross sectional area (CSA) of ACT EHM compared to controls (Supplementary figure 3A), we still found a significant DOX-dependent decrease in relative force of contraction in ACT EHM (by about 35%) after normalization to CSA (Figure 4E, F). Taken together, EHM from ACT iPSC CM showed an increased DOX-dependent beating frequency and a decreased DOX-dependent relative force generation compared to control iPSC CM.

**Figure 4.**
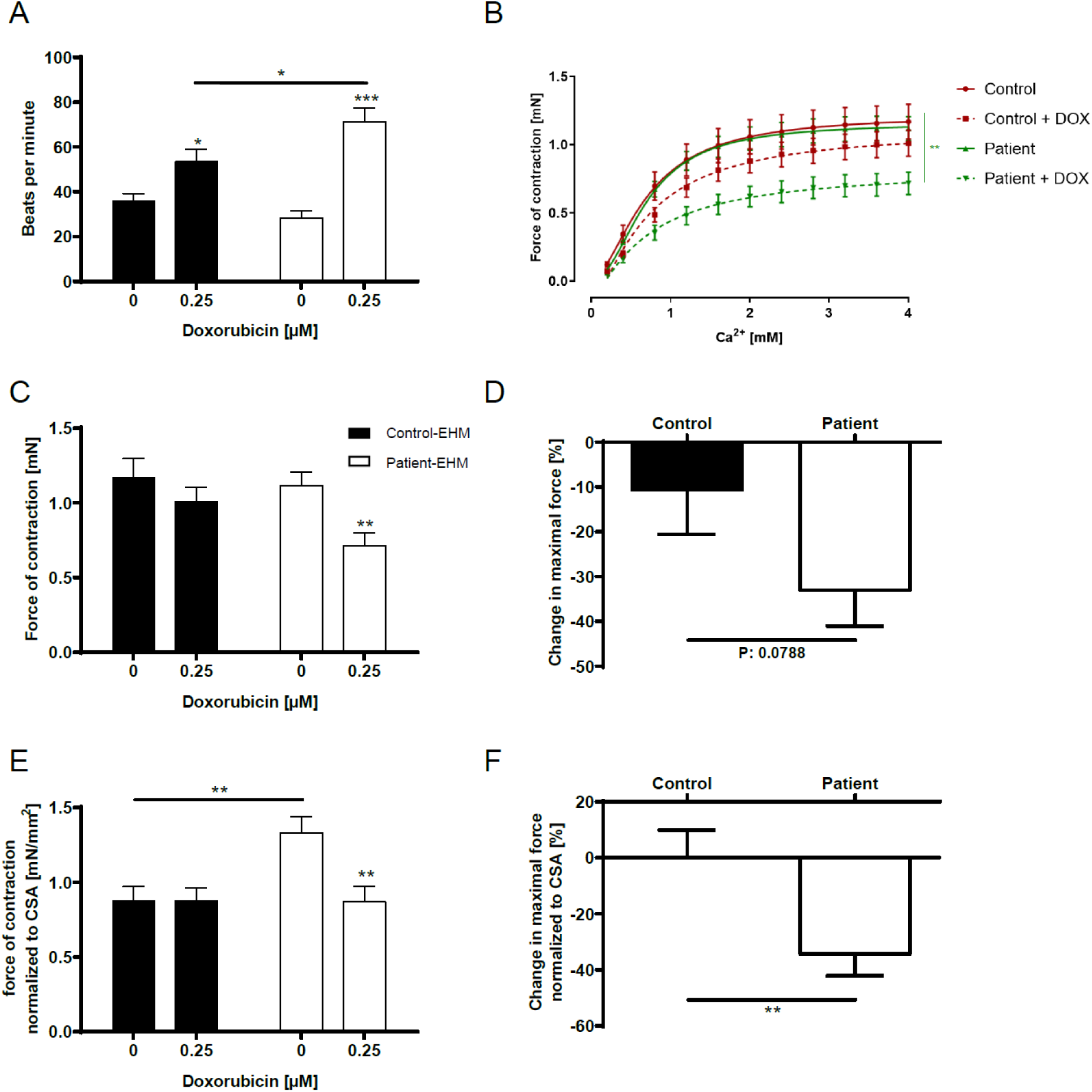
DOX decreases the force of contraction in engineered heart muscle (EHM). **(A)** Beating frequency of control and ACT patient EHM. **(B)** The generation of contractile force of control and ACT patient EHM is Ca^2+^-dependent, and impeded by DOX treatment. **(C)** Maximal force of contraction of control and ACT patient EHM is lowered after incubation with 0.25 µM DOX. **(D)** The DOX-induced relative change of maximal contractile force is greater in ACT patient EHM compared to control EHM. **(E)** Maximal force of contraction of control-EHM and ACT patient EHM normalized to cross sectional area (CSA). **(F)** DOX-induced change in maximal force of control and ACT patient EHM normalized to CSA. (**A**-**E**) Sample numbers: 12 control EHM from 2 differentiations, 18 ACT patient-EHM from 3 differentiations. Statistical analysis was performed using Student’s t-test or 2-way ANOVA analysis and Sidak’s multiple comparison test. Mean with SEM. * p < 0.05, ** p < 0.01, *** p < 0.001. * above single bars indicate statistical significances to untreated (0 µM DOX) conditions of the same group.

### Effect on doxorubicin on patient-specific gene expression

To further elucidate and confirm the molecular mechanisms underlying the pathogenesis of the DOX-induced phenotype, we performed RNA sequencing of EHM derived from three ACT patients and two controls, both with and without 0.25 µM DOX exposure for 24 h. We were able to identify the differentially regulated genes (DEGs) between the untreated EHM population and the EHM after DOX treatment (Figure 5A). After normalization for baseline expression, we identified 1106 upregulated DEGs and 990 downregulated DEGs in both ACT patient and control after DOX (Figure 5A). Principle component analysis (PCA) of the ACT and control groups treated and not treated with DOX showed a low variation within the groups, but allowed a clear distinction due to high variation between ACT and control as well as between -/+ DOX (Figure 5B). GO enrichment analysis of DOX-induced DEGs in both control and ACT-EHM identified significantly increased expression of genes with key functions including programmed cell death (10.8%, e.g. *BCL2L1, FAS, BAX, GPX1, ICAM1*), regulation of autophagy (e.g. *BECN1*), and catabolic processes (e.g. *STAT3, ABHD5*) (Figure 5C, Supplementary Figure 4A, Supplementary Table 1). Furthermore, we found significant decreases in actin cytoskeletal organization (18.03%, e.g. *ACTA2, MYBPC3, MYH6, MYH7, MYL2, MYL7, TNNI3, TNNT2*), muscle contraction (8.74%, e.g. *LDB3, MYBPC3, MYH6, MYL2, MYOM1, SORBS1, STMN1, TCAP, TNNT2*), and chromosome organization (2.19%, *BRCA2, TOP2A, TOP2B*) in both groups after DOX treatment (Figure 5D, Supplementary Figure 4B, and Supplementary Table 2). Normalized counts of DOX-induced regulated DEGs associated with general molecular mechanisms such as apoptosis (BAX, BCL2L2), catabolic processes (*ADHB5*) as well as with chromosome organization (TOP2A) are shown in Figure 5C and D. The downregulation of cardiac structural genes in response to DOX is illustrated in a heat map for control and ACT patients (Figure 5D), and was also established in animal *in vitro* and *in vivo* models(47) (48). We confirmed and validated a consistent downregulation of transcripts such as α-actinin *(ACTA2)*, α-myosin heavy chain *(MYH6)*, β-myosin heavy chain *(MYH7)*, and cardiac troponin T *(TNNT2)*, after DOX treatment by qPCR in human 2D iPSC CM in both populations (Figure 5E). These data suggest that DOX-treated EHM recapitulate general ACT disease mechanisms such as programmed cell death, myofibril damage or catabolic processes, and confirm the increased DOX-dependent apoptosis and sarcomeric dysregularity in iPSC CM of control and ACT patients (Figure 2D-H).

**Figure 5:**
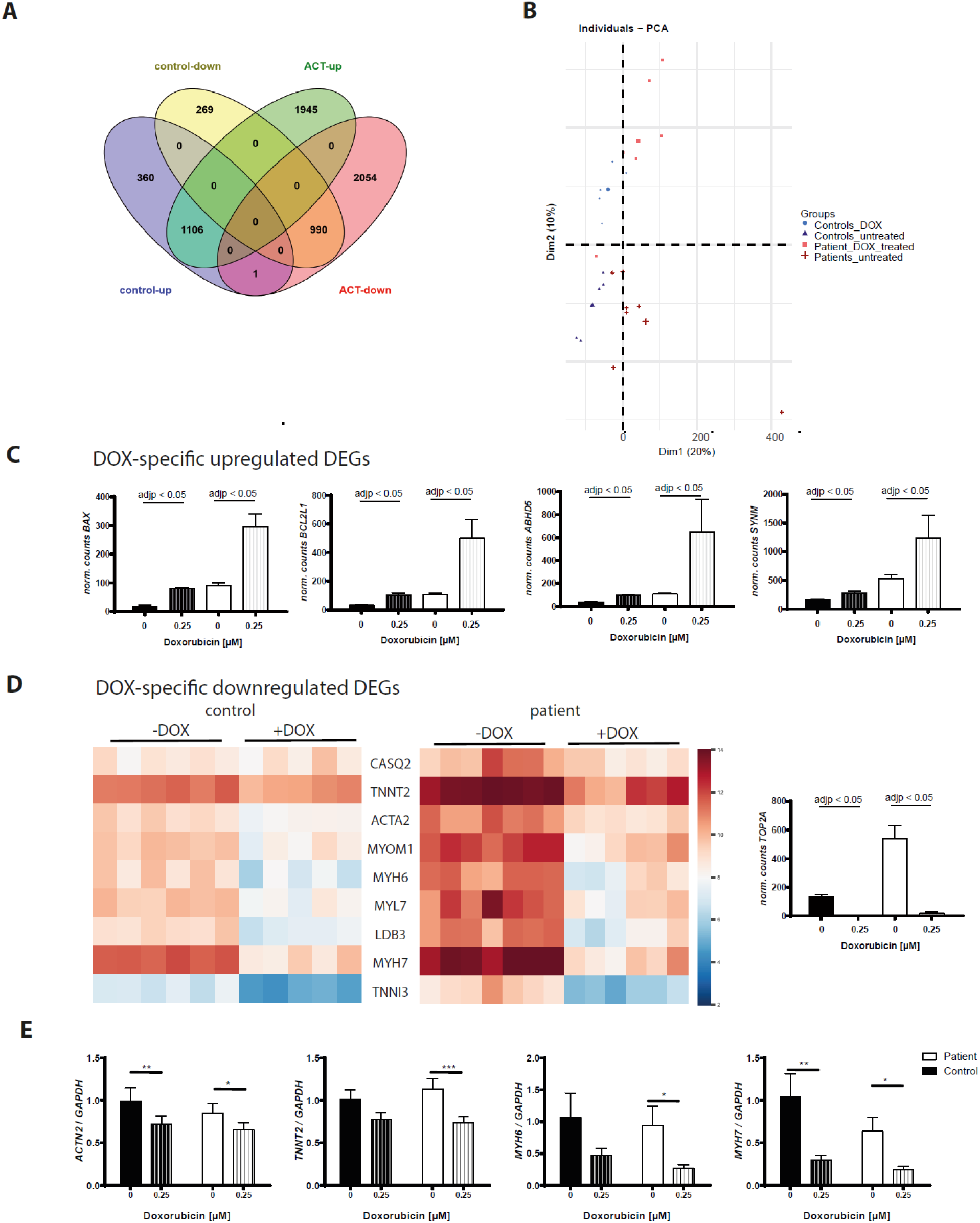
DOX-dependent differential gene expression in both ACT patient EHM and control EHM. **(A)** DOX-induced differentially expressed genes (DEGs) in ACT patient EHM and control EHM compared to untreated conditions illustrated in a Venn diagram. **(B)** PCA plot of control and ACT patient EHM samples used for the analysis. **(C)** Normalized counts of DOX-dependent upregulated DEGs (*BAX, BCL2L1, ABHD5, SYNM*) as well as downregulated DEG TOP2A (D) in both control (black) and ACT patient EHM (white). Adjp is based on analysis in A. **(D)** Heat map of 9 structural genes that are significantly downregulated after DOX treatment in control and ACT patient EHM. **(E)** Differential DOX-dependent downregulation of *ACTN2, TNNT2, MYH6*, and *MYH7* were confirmed by qRT-PCR in control as well as ACT patient iPSC CM. Mean with SEM. *P < 0.05; **P < 0.01; ***P< 0.001; untreated versus treated by Student’s t-test.

Next, we focused on signaling pathways which are specifically regulated in control EHM, but not in ACT EHM, after DOX treatment. Interestingly, we found 360 upregulated and 269 downregulated genes which were specifically altered in expression in control EHM after DOX (Figure 5A). GO-term analysis of control DOX-induced DEGs identified significantly increased expression of translation, translational initiation (*EIF3D, EIF6*), ribosome assembly (*EFNA1*), and positive regulation of p38MAPK cascade (*PRMT1*) (Figure 6A, Supplementary Table 3). Normalized counts of control-specific DOX-dependent DEGs were associated with increased translation as well as p38MAPK signaling, as depicted in Figure 6A.

**Figure 6.**
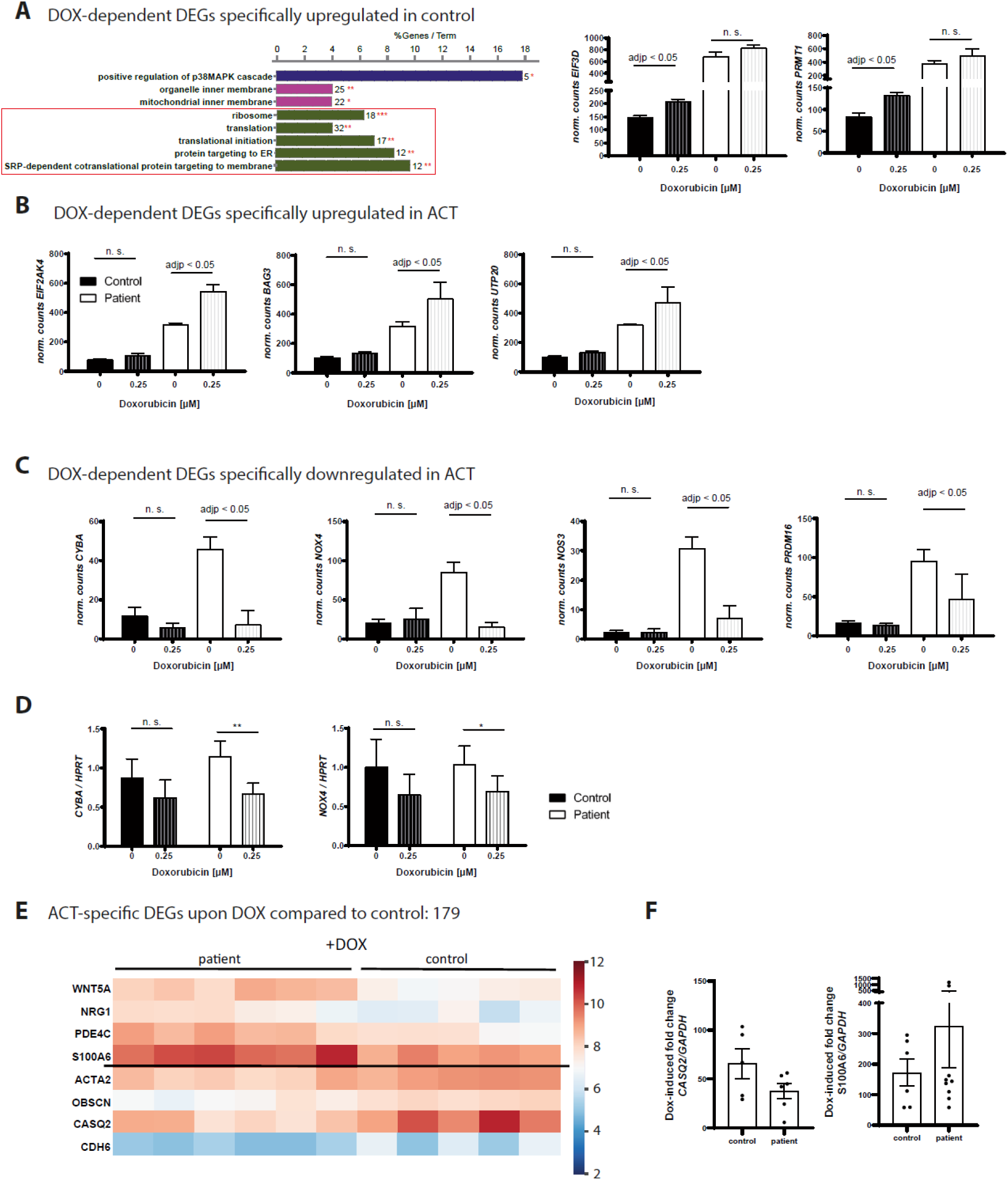
DOX-dependent modulation of gene expression in EHM of ACT patients and controls. **(A)** Significant enriched GO terms after DOX in control EHMs compared to untreated conditions according to ClueGo Cytoscape plugin. The data of differential expression are based on all up-regulated genes. Data are generated by GO cellular component, GO biological process or KEGG pathway analysis. Terms of same color correspond to terms containing a similar group of genes. The bars represent the number of the genes from the analyzed cluster that were found to be associated with the term, and the label displayed on the bars is the percentage of genes found compared to all the genes associated with the term. *P < 0.1; **P < 0.05 calculated based on hypergeometric distribution from Database for Annotation, Visualization and Integrated Discovery (DAVID, v6.7). Normalized counts of control-specific, DOX-dependent upregulated DEGs (*EIF3D, PRMT1*). **(B, C)** Normalized counts of ACT-specific, DOX-dependent upregulated DEGs (*EIF2AK4, BAG3, UPT20*) as well as downregulated DEGs (*CYBA, NOX4, NOS3, PRDM16)*. Adjp is based on analysis in A. **(D)** Differential DOX-dependent downregulation of NADPH-oxidase subunits *NOX4* and *CYBA* were confirmed by qRT-PCR in ACT-iPSC-CM. Mean with SEM. *P < 0.05; **P < 0.01 by Student’s t-test. **(E, F)** DEGs specifically regulated between control and ACT patient EHM after DOX. **(E)** Heat map of 8 important genes that were differentially expressed between control and ACT EHM after DOX treatment. **(F)** Differential DOX-dependent higher downregulation of *CASQ2* and higher upregulation of *S100A6* in ACT EHM compared to control EHM was confirmed by qRT-PCR in ACT and control iPSC-CM. Mean ± SEM. *P < 0.05; by Student’s t-test. For control EHM, 5-6 independent EHM from 1 cell line of control patient 1 and 2, and for ACT patient EHM, 6-7 independent EHM from 1 cell line of each of the 3 ACT patients (ACT1, ACT2, ACT3) were analyzed.

In contrast to control EHM, we found 1945 upregulated and 2054 downregulated genes in ACT patient EHM after DOX treatment (Figure 6A). The upregulated genes were involved in negative regulation of protein synthesis (Eukaryotic Translation Initiation Factor 2 Alpha Kinase 4 (EIF2AK4)) or protein nuclear export/ incorrect folding (BAG3) (Supplementary Figure 4C, Figure 6B, Supplementary Table 4). In addition, we found a significant decrease in oxidative stress-related genes (*CYBA, NOX4, NOS3*) in ACT EHM after DOX compared to the untreated patient group (Supplementary Figure 4C, Figure 6C, Supplementary Table 5). Of note, PR domain containing 16 (PRDM16), a repressor of TGFβ signaling and known genetic cause of left ventricular, non-compaction cardiomyopathy (LVNC) and DCM (49), was significantly downregulated in ACT iPSC CM after DOX, but not in DOX-treated control cells (Figure 6C). Normalized counts of ACT-specific DOX-dependent DEGs associated with translation (*EIF2AK4, BAG3*) as well as redox stress (*CYBA, NOX4, NOS3*), or *PRDM16* are shown in Figure 6B, C. We validated the ACT-specific downregulation of redox stress-associated genes *CYBA* and *NOX4* in response to DOX by qPCR in 2D iPSC CM (Figure 6D). These data indicate differentially regulated mechanisms such as mRNA translational processes in DOX-treated ACT EHM compared to control.

In order to identify different impacts of DOX in ACT and control EHM, we detected 179 significantly regulated genes with 71 downregulated and 108 upregulated genes after DOX treatment in ACT and control patients (Supplementary Table 6, Figure 6E, F). We were able to show that cardiac structural genes such as *ACTA2* or *OBSCN* are significantly more downregulated in response to DOX in ACT iPSC CM compared to control cells as shown in a heatmap (Figure 6E). Furthermore, calsequestrin *(CASQ2)*, which is involved in calcium binding in muscle cells, was significantly downregulated after DOX treatment in ACT patients as compared to controls. In addition, differences in other calcium handling-related genes such as the calcium sensor S100, the calcium-binding protein S6 (*S100A6*), or the calcium binding-related gene cadherin 6 (*CDH6*) were identified in this group (Figure 6D). We confirmed and validated the increased downregulation of *CASQ2* as well as the increased upregulation of S100A6 after DOX treatment in ACT EHM by qPCR in 2D iPS -CM (Figure 6F). The different DOX-dependent regulation of genes such as WNT5A, NRG1 and PDE4C in ACT and control patients points to the involvement of several more signaling pathways in ACT.

### Disturbed functionality of calcium homeostasis in patients with ACT after DOX

Due to DOX-dependent alterations in cardiac functionality and DEGs in ACT EHM, we sought to investigate calcium kinetics in iPSC CMs at a therapeutic DOX dose of 0.25 µM as well as a toxic DOX dose of 5 µM. We detected a significantly higher T50 transient decay time in ACT-iPSC CMs compared to controls at DOX concentrations of 0.25 µM and 5 µM (Figure 7A). Furthermore, we found a decrease in the amplitude of calcium transients (Ca2^+^_i_) in the ACT iPSC CMs as compared with control subjects at DOX concentrations of 0.25 µmol/l (Figure 7B), but no difference between the groups at DOX concentrations of 5 µmol/l (Figure 7B). While our results showed a higher rise time in ACT-iPSC CMs compared to controls at low DOX concentrations, 5 µmol/l DOX did not result in a change of this parameter between the groups (Figure 7C).

**Figure 7.**
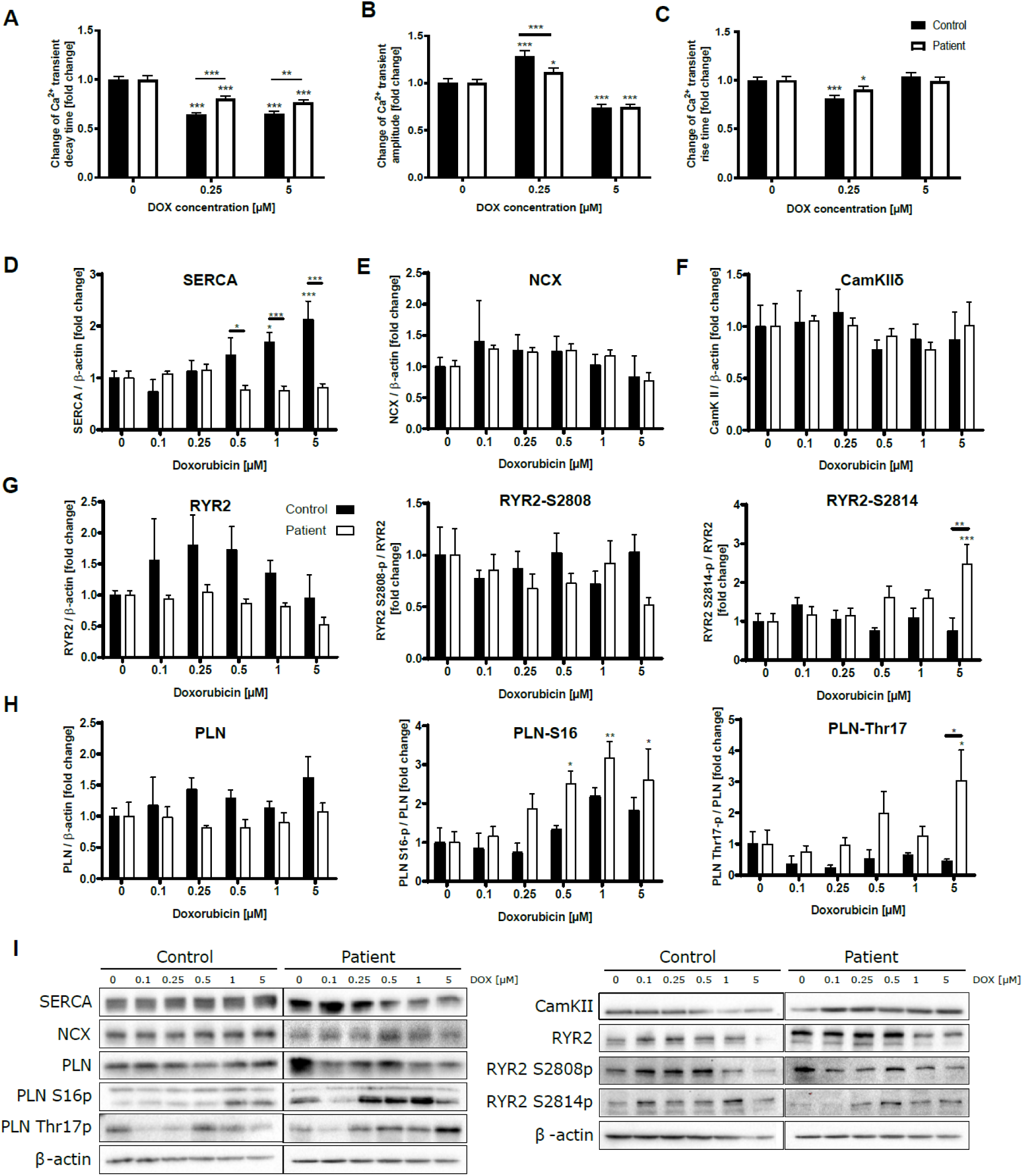
DOX disturbs Ca^2+^ homeostasis. (**A-C**) Effects of DOX (0.25 µM and 5 µM) on Ca^2+^ transient decay time, Ca^2+^ transient amplitude, and Ca^2+^ transient rise time in iPSC-CM of control and ACT patients. **(D-H)** DOX-induced relative changes in the amount of protein expression and phosphorylation of SERCA, NCX, CamKIIδ, RYR2, RYR2-S2808, RYR2-S2814, PLN, PLN-S16p, PLN-Thr17p in iPSC-CM of control and ACT patients after 24 h of DOX treatment. (**I**) Representative Western blots of calcium-associated proteins in DOX-treated iPSC CM of control and ACT patients. Sample numbers: **(A-C)** 200-61 control-iPSC-CM from 9 differentiations, 197-119 ACT patient-iPSC-CM from 10 differentiations. **(D-H)** 4 control-iPSC-CM differentiations, 6 ACT patient iPSC CM differentiations. Statistical analysis was performed using 2-way ANOVA with Dunnett’s multiple comparison test. Mean with SEM. * p < 0.05, ** p < 0.01, *** p < 0.001. * above single bars indicate statistical significances to untreated (0 µM DOX) conditions of the same group.

To better understand the impact of DOX on calcium kinetics, key calcium regulating proteins and their phosphorylation were analyzed using Western blots. Interestingly, in controls, iPSC CM expression of SERCA increased in a DOX dose-dependent manner (0.5 – 5 µmol/l), whereas SERCA expression decreased in ACT-iPSC CM at the same DOX concentrations (Figure 7D and I). Additionally, we detected an increased expression of ryanodin receptor 2 (RYR2) upon DOX treatment in control cells, but not in ACT-iPSC CM (Figure 6G, I). In contrast, the expression of sodium-calcium exchanger 1 (NCX1) and phospholamban (PLN) upon DOX treatment was not affected in either group (Figure 7E, H and I). However, PKA-specific phosphorylation of PLN-S16 increased significantly in a DOX dose-dependent manner, predominantly in ACT-iPSC CM (Figure 7H, I). In contrast, PKA-specific phosphorylation of RYR-S2808 did not result in DOX-induced changes (Figure 7G). The amount of calcium/calmodulin (CaM)-dependent kinase II δ (CamKIIδ) was not regulated after DOX treatment in both groups (Figure 7F). Of note, CamKIIδ-specific phosphorylation of RYR2-S2814 or PLN-Thr17 increased in ACT-iPSC CM depending on the dosage of DOX, with a significant increase at 5 µM DOX compared to basal conditions and control-iPSC CM for both targets (Figure 7G, H, I). We also quantified the basal protein expression of calcium-handling proteins, and detected no significant differences in the amounts of SERCA, NCX1, PLN, CamKIIδ, and RYR2. In addition, the phosphorylation of PLN-S16 or -Thr17 and RYR2-S2808 or - S2814 were similar under basal conditions in control and ACT-iPSC CM (Supplementary Figure 5A, B).

In summary, these results suggest that there are differing Ca^2+^ mechanims in the control and ACT groups that react to acute DOX treatment, triggering e.g. an increase in RYR2 depending on DOX dosage. SERCA mRNA translation was for example only observed in control iPSC CM, and CaMKII-dependent phosphorylation of important calcium-handling proteins was increased in ACT iPSC CM depending on DOX dosage; This CaMKII-mediated phosphorylation is probably ROS-dependent and a compensatory reaction to acute DOX since translational initiation processes are disturbed in ACT iPSC CMs.

### Genetic-based cardiac dysfunction in patients with ACT

The high inter-individual variance in ACT manifestation might be due to genetic predisposition. To test this, we performed whole exome sequencing (WES) on DNA from skin fibroblasts of three ACT patients and two controls used in this study for the iPSC generation. Based on the hypothesis of a genetic predisposing variant with a strong functional effect, we focused our WES data analysis on the 81 genes previously associated with various genetic forms of cardiomyopathies and cardiac arrhythmias. We found that patient ACT-2 carried the heterozygous c.C2633T variant in *PRDM16* (p.P878L) that was predicted to be deleterious or damaging by different computational prediction programs. This variant does not represent a common single nucleotide polymorphism (SNP), and is not listed in over 121,000 alleles in the ExAC database. Causative variants in *PRDM16* are associated with LVNC and DCM (49), and therefore the PRDM16 p.C2633T variant from our study might be associated with ACT subject cardiotoxicity. Patient ACT-3 was found to carry the heterozygously rare variant c.3010G>A (present in 2 of <121,000 in the ExAC) in the Synemin (*SYNM*), which is predicted to substitute valine at position 1004 against isoleucine, and to cause damage. No likely candidate variant was found in the other ACT-1 patient. We confirmed all candidates by Sanger sequencing (data not shown).

## Discussion

In the present study, we showed for the first time that human iPSC CM and EHM can recapitulate a B cell lymphoma patient’s predilection to DOX-induced cardiac dysfunction when exposed to clinically high levels of DOX. ACT-iPSC CM derived from lymphoma cancer patients are consistently more sensitive to DOX toxicity and demonstrate higher intracellular DOX amounts after treatment, disorganized myofilament structure as well as increased cell death. An increase in DOX-dependent ROS production was not only identified in ACT-iPSC CM, but also in ACT cFB that were isolated from strong fibrotic human left ventricular ACT myocardium. We were also able to demonstrate that ACT-iPSC CM are not able to adapt to acute DOX stress by modulating expression or activity of important Ca^2+^ handling proteins such as SERCA and RYR2, as was the case in control iPSC-CM. The molecular and structural differences observed here may be the result of DOX-dependent, impaired mechanical functionality in ACT EHM. Furthermore, genetic variants of several key regulators of cardiac function were identified, suggesting a genetic predisposition in ACT. The differences in the responses of control and ACT cFB, iPSC CM and EHM to DOX are in line with those observed in ACT patient myocardium and in animal models with DOX-induced ACT (29). This confirms their suitability as model systems for studying the predilection of high-risk patients and the molecular and pathophysiological mechanisms underlying DOX-dependent cardiac dysfunction.

Our study adds fundamental knowledge to recent reports demonstrating the use of iPSCs from breast cancer patients to recapitulate their predilection to ACT (40, 41). To our knowledge, ours is the first human iPSC CM model of DOX-induced cardiac dysfunction in patients with aggressive B cell lymphoma who were treated with high doses of DOX. In our model, the ACT phenotype can not only be studied in a 2D monolayer of iPSC CM, but in a myocardium-like, more mature human 3D culture(50), allowing for a functional and morphological comparison to the myocardium of end-stage heart failure ACT patients. Furthermore, we were able to reveal on a patient-specific level the activity and expression of important Ca^2+^ handling proteins such as SERCA, RYR2 and CamKIIδ. Our focus was on both ACT iPSC CM as well as ACT cFB as a non-CM cell type, and how both contribute to ACT development after DOX treatment.

Reduced ventricular ejection fraction and myocardial contractile dysfunction are characteristic of patients with DOX-induced cardiac dysfunction as well as with dilated cardiomyopathy (51). In this study, we analyzed the DOX-dependent ACT phenotype and function in a 3D myocardium of patient-specific iPSC CM (50), which resembles the in vivo heart. We demonstrated a decreased force of contraction in DOX-treated ACT EHM, similar to clinical features of ACT patients and to iPSC EHM of patients with DCM (39). In addition, we demonstrated a DOX-dependent increased beating frequency in ACT EHM that may mimic potential rhythm disturbances in patients with acute ACT immediately after DOX treatment (51).

Interestingly, in monolayer iPSC-CM we observed multiple DOX-associated processes relevant to ACT such as a dose-dependent increase in extracellular H2O2, apoptosis, and sarcomeric disarray, which were significantly more pronounced in ACT-iPSC CM compared to controls. These processes are supported by transcriptome analysis of ACT and control EHM of this study, revealing significant general DOX-dependent gene expression changes in pathways related to “programmed cell death”, “ROS production”, or “striated muscle contraction” in both groups. This is in line with recently published studies showing general DOX-dependent changes in gene expression (41, 52). Of note, DOX concentrations greater than 0.5 µM (0.75 - 5 µM) were not associated with increased H2O2 amounts but rather higher apoptosis in both ACT and control iPSC-CM. This indicates that therapeutic DOX doses (<0.5 µM) may reflect processes resembling the phenotype of delayed chronic cardiotoxicity, while toxic doses (> 0.5 µM) may resemble acute cardiotoxicity (52).

Studies on DOX concentrations in plasma during treatment suggest rapid clearance of the drug from circulation (53). Here we report for the first time that DOX levels were close to the detection limit after 7 days of recovery from one 24h DOX interaction, whereas DOX-dependent increase in H2O2 was detectable over weeks. Our results provide proof for the assumption that initial DOX molecules do not remain in cardiac cells for extended periods, but cause impairments that lead to long-term cardiotoxicity in a hit-and-run manner (54).

Another important finding includes the altered response to calcium kinetics observed in iPSC CMs of patients with ACT after DOX treatment. Due to decreased cardiac force of contraction in DOX-treated ACT EHM, we found lower SERCA and RYR2 protein expression (transcriptome data confirmed a DOX-dependent decreased expression of RYR2 only in the ACT group) in ACT iPSC CM compared to control iPSC CM. This is consistent with a higher absolute T50 time in ACT iPSC CMs and a decrease in the amplitude of calcium transients in ACT iPSC CMs compared to controls under the same conditions. These data indicate slower re-uptake of calcium into SR vesicles by the SERCA-PLN complex and retarded relaxation in ACT compared to control. This is in line with the frequently reported decreased DOX-dependent or independent SERCA2a activity in heart failure myocardium in the present (Figure 1D) and in previous studies (30, 55), which is functionally manifested in lower Ca^2+^ uptake into the SR and cytoplasmic Ca^2+^ overload. Furthermore, SERCA was shown to be inhibited in its activity by high DOX levels (18). In addition, our data on increased CamKIIδ activity in ACT iPSC-CM compared to control may be due to increased DOX-dependent ROS production along with greater oxidation and thereby activation of CamKIIδ in ACT, as has been shown in previous studies (29). Inhibition of CaMKIIδ improves dysfunction of a failing heart (56). Interestingly, transcriptome analysis revealed that protein arginine methyltransferase 1 (PRMT1), which is essential for preventing cardiac CaMKII hyperactivation, is specifically upregulated in control EHM upon DOX treatment, but not in the ACT EHM. PRMT1 methylates CaMKII and thereby inhibits its activity, thus protecting against pathological responses (57). Of note, biphasic dose-dependent effects of DOX were shown for RYR2 expression in control iPSC CMs with increased amounts at 0.25 µM DOX and similar levels at 5 µM at baseline; this difference in DOX-dependent expression of RYR2 is probably a function of the increased Ca transient amplitude at low DOX in iPSC CM as compared to those at high DOX concentrations. Similar results were shown in previous studies, where a reversible DOX-dependent RYR2 activation was described at low DOX concentrations and a DOX-mediated inhibition of RYR2 at high DOX concentrations such as 5 µM (18).

In conclusion, this study provides new evidence that different Ca^2+^ regulatory mechanisms play major roles in ACT-iPSC CM as opposed to control CM. Control iPSC CM were able to adapt to acute DOX stress by increasing protein expression of important calcium regulatory proteins such as RYR2, SERCA, or PRMT1. In contrast, ACT-iPSC CM were not able to react to acute DOX stress by mRNA translational processes, and showed increased post-translational processes such as CamKIIδ-mediated phosphorylation. These novel data point to distinct mechanisms that may explain the different severities of DOX-dependent effects on Ca^2+^ functional processes in ACT and controls. Of note, we found a DOX-dependent downregulation of *CASQ2* in ACT iPSC CM compared to control CM, which may have resulted in reduced Ca^2+^ binding capacity of CASQ2, along with a reduced SR Ca^2+^ release and altered SR Ca^2+^ storage capacity. This would be in line with previous studies showing that anthracyclines bind with a high affinity to cardiac CASQ2, and thereby reduce the Ca^2+^ binding capacity of CASQ2 (58-60). Nevertheless, detailed functional Ca^2+^ measurements, including comprehensive analysis of SR Ca^2+^ content, Ca^2+^ leak and SERCA function, will be necessary in future studies.

Our hypothesis that DOX treatment could have different impacts on mRNA translation in ACT and controls CM is strengthened by transcriptome data showing DOX-induced DEGs with increased expression of translation, translational initiation (EIF3D, EIF6), and ribosome assembly (EFNA1) in control EHM. In contrast, DOX-treated ACT EHM demonstrated upregulated DEGs, including negative regulation of mRNA translation-associated genes such as the metabolic-stress sensing protein kinase EIF2AK4. EIF2AK4 is known to phosphorylate EIF2α as a competitor for translational initiation, leading to repression of global protein synthesis (61). It was previously reported that the translation was reduced by only 2% in human keratinocytes (62) and 75% in prostate cancer cells (63). White and colleagues (2007) demonstrated that decreased translation was caused by sustained phosphorylation of elongation factor 2 (EF-2). Phosphorylation of EF-2 occurred in a kinase-independent manner, most likely through elevated ROS levels (63).

Our observation that ACT patient-specific iPSC-CM and cFB displayed a more severe cardiac phenotype following DOX treatment than the control groups indicates that ACT patients may possess combinations of genetic variants associated with increased susceptibility to DOX-induced cardiac dysfunction. For that reason, we performed WES and identified highly relevant variants in cardiac genes, such as PRDM16 (c.C2633T) or SYNM (c.3010G>A; pV1004I) in patients ACT 2 or 3. Interestingly, autosomal dominant mutations in PRDM16 have been associated with LVNC and DCM (OMIM 615373). Sequence analyses revealed that the identified PRDM16 variant substitutes the highly conserved proline at position 878 with leucine (p.P878L). It was previously shown that single variants or a human truncation mutant in zebrafish resulted in contractile dysfunction and impaired cardiomyocyte proliferation capacity (49). These findings further support the suggested functional evidence of variants in PRDM16 in ACT. Synemin is an intermediate filament protein and its absence in mice was demonstrated to cause profound structural and functional abnormalities in the heart (64). Previously, a novel heterozygous variant p.(Trp538Arg) of SYNM was identified in three generations of a family with four patients with DCM (65). The exact molecular function and the effect of the identified novel PRDM16 and SYNM variants on the ACT phenotype need to be analyzed in follow-up studies. Of note, *PRDM16* and *SYNM* are both differentially expressed in the EHM transcriptome data of this study. While *PRDM16* is significantly decreased specifically in ACT-EHM after DOX (Figure 9B), *SYNM* seems to be a general DOX target since it is significantly upregulated in both ACT and control after DOX (Fig. 8C). To prove the hypothesis of a genetic predisposition to ACT, further WES experiments in larger patient cohorts will be necessary. Nevertheless, in 2 of 3 patients we identified variants in important cardiac proteins, which are associated with cardiac pathologies.

In this study we focused on several cardiac cell types including ACT cFB as non-CM cell type, isolated from human ACT myocardium, which were suggested to contribute to ACT by amplifying development of fibrosis upon DOX application (66). We observed that ACT cFB show specific cFB morphology, proliferation, and marker expression. DOX-treated ACT-cFB markedly increased expression of *ACTA2* and *COL1A1*, indicating an increased myofibroblastic stress response. In addition, SERCA was increased after DOX, suggesting that the Ca^2+^ response in cFB is a result of pathological stimuli, as was shown in previous studies (67). However, Ca^2+^ signaling in cardiac fibroblasts is still not fully understood (67). ACT-cFB produced more ROS upon DOX compared to commercially obtained healthy cFB as well as sFB, suggesting that ACT-cFB are more sensitive to anthracyclines than the other two FB types. The DOX-dependent increase in ROS may serve as the trigger for cFB activation, ECM damage and contribution to ACT development. This is in line with DOX-dependent effects in human cFB, inducing trans-differentiation of cFB to myofibroblasts at low concentrations in an organ- and species-dependent manner(68). Whether the response of ACT-cFB to injury by proliferating in or migrating to areas of damage was influenced by DOX treatment in our study needs to be elucidated in the future.

It is also important to point to the limitations of our study. First, we used two different iPS cell lines per ACT patient and control, all generated by non-integrative methods, for a minimum of 3 cardiac differentiation experiments. The maturity of iPSC CM as well as the patient’s genetic background, age and sex may have influenced the results. In future studies, age-matched control iPSCs without previous DOX treatment should serve as ideal controls. Secondly, as the functional consequences of the failed adaptation to DOX stress in ACT-iPSC-CM by modulating expression or activity of Ca^2+^ handling proteins (SERCA and RYR2) is still unclear, detailed functional Ca^2+^ measurements, including comprehensive analysis of SR Ca^2+^ content, Ca^2+^ leak and SERCA function, will required in the future. Thirdly, our data suggest the contribution of novel *SYNM* and *PRDM16* variants to the observed ACT phenotype, but the mechanisms are unclear. Therefore, detailed functional analyses including genome editing work needs to be done in the future. Finally, aside from the studied DOX effects, further analysis of potential cardio-protective mechanisms and therapeutic substances, including ranolazine and dexrazoxan, need to be investigated.

In conclusion, we used cFB, iPSC CM and EHM from healthy controls and ACT patients with cardiac dysfunction to demonstrate that all cell types can recapitulate a patient’s predilection to ACT after exposure to DOX. The iPS CM showed characteristics comparable to mechanical dysfunctional and strong fibrotic human cardiac tissue from ACT patients with end-stage heart failure. Our results suggest that altered Ca^2+^ signaling and translational processes underlie CM dysfunction. Targeting of the Ca^2+^ signaling pathway or translation-associated components may thus represent a new approach for mitigating the cardiac side effects of DOX in cancer patients. Genetic variants associated with ACT development could also provide new targets for therapy of ACT.

## Material & Methods

Details are available in the Supplementary Appendix.

### Human myocardium of ACT patients

We obtained left ventricular tissues from explanted hearts of 5 ACT patients with end-stage HF who were undergoing heart transplantation (EF ≤ 20%). After explantation, the heart was immediately placed in pre-cooled cardioplegic solution containing NaCl (110 mM), KCl (16 mM), MgCl2 (16 mM), NaHCO3 (16 mM), CaCl2 (1.2 mM), and glucose (11 mM). Myocardial samples for qPCR, Western blot and immunocytochemistry stainings (fibrosis) were immediately frozen in liquid nitrogen and stored at −80°C. We used control myocardium from healthy donor hearts that could not be transplanted for technical reasons for qPCR, immunoblots and fibrotic stainings. Collection of human samples was in compliance with the local ethical committee (Az. 31/9/00), and written informed consent was received from each participant before transplantation.

### Selection of ACT patients for iPSC generation

Patients described in this study were initially identified as lymphoma patients and selected from the ‘rituximab with CHOP over age 60 years’ (RICOVER60) trial (NCT00052936). The study cohorts comprised 61-80 year-old patients with aggressive CD20+ B cell lymphomas treated with six or eight cycles of CHOP-14 with or without rituximab. All patients of this study had been treated with doxorubicin (DOX) as part of a CHOP treatment. ACT patients 1-3 suffered from anthracycline-induced cardiotoxicity, whereas control patients did not develop cardiac symptoms.

### Human skin and cardiac fibroblast isolation and cultivation

The study was approved by the ethical committee of the University Medical Center Göttingen (Az 21/1/11 and 31/9/00). The fibroblast culture was derived from skin punch biopsies of the donors (RICOVER60 study) or from human cardiac tissue (approximately 1 mm^3^) from the left ventricle of ACT end-stage heart failure patients. Somatic cells were isolated as previously described by our group (38). To secure attachment and proliferation of the fibroblasts, the dish was left without medium change for one week, after which the tissue was removed. Primary skin or cardiac fibroblasts were cultured in fibroblast growth medium composed of DMEM supplemented with 10% FCS, 1x NEAA, glutamine (2 mmol/L), β-ME (50 µmol/L), penicillin (50 U/mL)/streptomycin (50 µg/mL), and bFGF (10 ng/mL) at 37°C with 5% CO2 atmosphere. The medium was replaced every second day and cells were passaged once to twice per week. FB were used from passage 3-6.

## Author contributions

LH generated and characterized the iPSCs, performed experiments (sarcomeric integrity, apoptosis, ROS, calcium), and contributed to manuscript writing. AM performed experiments (DOX incorporation, ROS), and contributed to manuscript writing. MT performed and analyzed the EHM experiments and received funding. SK generated the expression data of cardiac tissues. SK and WB generated the cardiac fibroblast data. SS provided healthy and diseased human cardiac tissue, contributed to concept development and received funding. AS and GB performed the quantification of fibrosis in human cardiac tissue slices. CXS and AMS provided guidance during DOX incorporation experiments. RD performed the in vivo differentiation experiments. LW provided patient data. JH, RT and BM generated the RNAseq data of the EHM and contributed to analysis. BM received funding. YL and BW performed the WES data and analysis. GH contributed to study design. KSB developed the concept, designed the study, analyzed the RNAseq data, received funding and wrote the manuscript.

## Conflict of interest

The authors share no conflict of interest

### Acknowledgments

The authors thank Sandra Georgi, Carmen Klopper, Johanna Heine, Yvonne Metz, and Timo Schulte for excellent technical assistance; Dr. Alexander Becker and all others who helped collect skin punch biopsies from patients with ACT and control subjects, and Hanibal Bohnenberger for organizing hematoxylin and eosin staining of human cardiac tissue slices in the Department of Pathology, University Medicine Göttingen. We thank Ms Marita Ziepert for providing the information on the ACT patients from the RICOVER60 study as prerequisite for choosing the ACT patient in this study. In addition, we thank Lydia Maus and Cynthia Bunke for proofreading.

## Funding

This work was supported by the Bundesministerium für Bildung und Forschung (BMBF) grant e:Bio – Modul II – Verbundprojekt: CaRNAtion [031L0075C to KSB and GH], the German Center for Cardiovascular Research (DZHK) [B14031KSB to KSB, MT, and BM], and the Heidenreich von Siebold Program from the University Medical Center Göttingen (KSB), the German Heart Foundation/German Foundation of Heart Research [AZ. F/38/18] (to KSB), and the Else Kröner-Fresenius-Stiftung Foundation [2017-A137] (to KSB and STS). Andreas Maus, Steffen Köhne, and Wiebke Maurer are fellows of the International Research Training Group 1816 funded by the Deutsche Forschungsgemeinschaft. AMS is supported by the British Heart Foundation and in part by the Department of Health via a National Institute for Health Research (NIHR) Biomedical Research Centre award to Guy’s & St Thomas’ NHS Foundation Trust in partnership with King’s College London.

## Central Figure

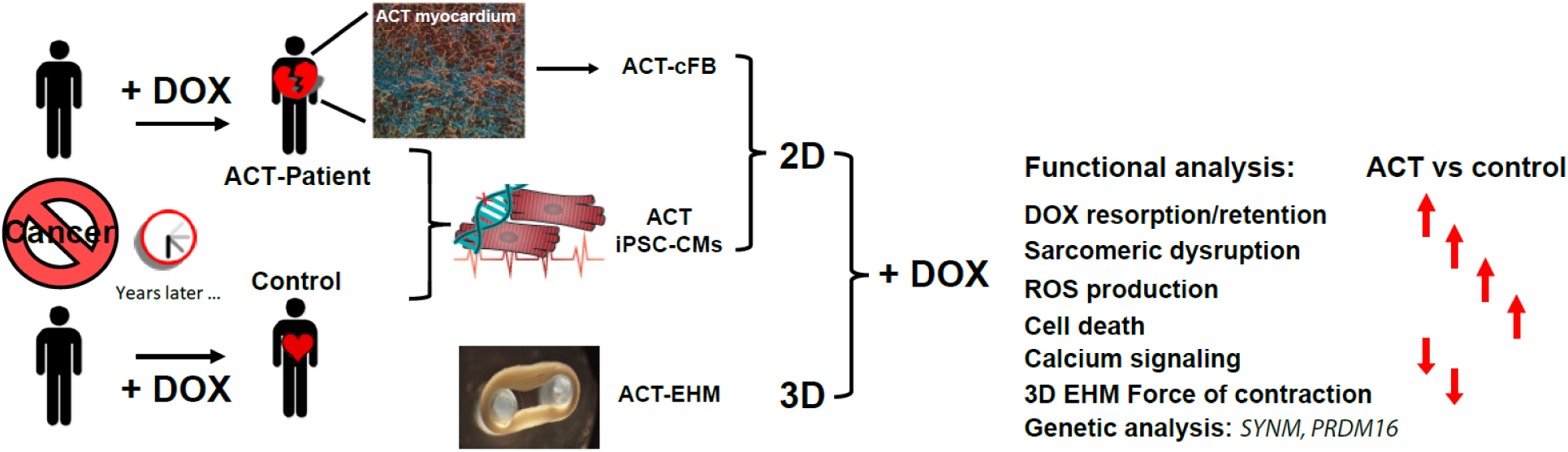

